# *Loss of miR-9-2* Causes Cerebral Hemorrhage and Hydrocephalus by Widespread Disruption of Cell-Type-Specific Neurodevelopmental Gene Networks

**DOI:** 10.1101/2025.07.31.668014

**Authors:** SP Fregoso, M Atapattu, LK Callies, D Monet, A Leonardson, L Clark, S Xu, TJ Cherry

## Abstract

*MIR-9-2* is a broadly and highly expressed microRNA in the developing brain and is frequently deleted in 5q14.3 Microdeletion Syndrome, a rare but severe neurodevelopmental disorder. Despite this, little attention has been paid to the unique contributions of *MIR-9-2* to neurodevelopment and disease. We find that deletion of this microRNA leads to embryonic cerebral hemorrhages and severe hydrocephalus, while disrupting gene networks across a wide range of cell types in the developing brain, thus revealing underappreciated and non-redundant molecular, cellular, and system-wide functions for *MIR-9-2* in neurodevelopment.

## Main

MicroRNAs are post-transcriptional repressors of target mRNA translation and play crucial roles in neurodevelopment^1^. The miR-9 family of microRNAs is highly and widely expressed during Central Nervous System (CNS) development and is known to regulate neural proliferation, differentiation, and maturation in many species^2,3^. In humans, one family member, *MIR-9-2*, has been associated with autism spectrum disorder risk^4,5^ and is frequently lost in 5q14.3 Microdeletion Syndrome^6-8^ (Fig. 1A), a neurodevelopmental disorder characterized by developmental delay, disrupted or absent speech, epilepsy, and frequently by cerebral vascular malformations^9,10^ and enlarged cerebral ventricles ^9,11^. The features of 5q14.3 Microdeletion Syndrome are typically attributed to loss of protein-coding genes at this locus including *MEF2C*^6,12-14^, *TMEM161B*^15^, or *RASA1*^10,12^; however, the *MIR-9-2HG* (host gene) is more highly and broadly expressed across cell types in the developing human brain^16^ (Fig. 1B), indicating that this specific MIR-9 family member may be an overlooked and significant contributor to this human disorder.

**Fig. 1.**
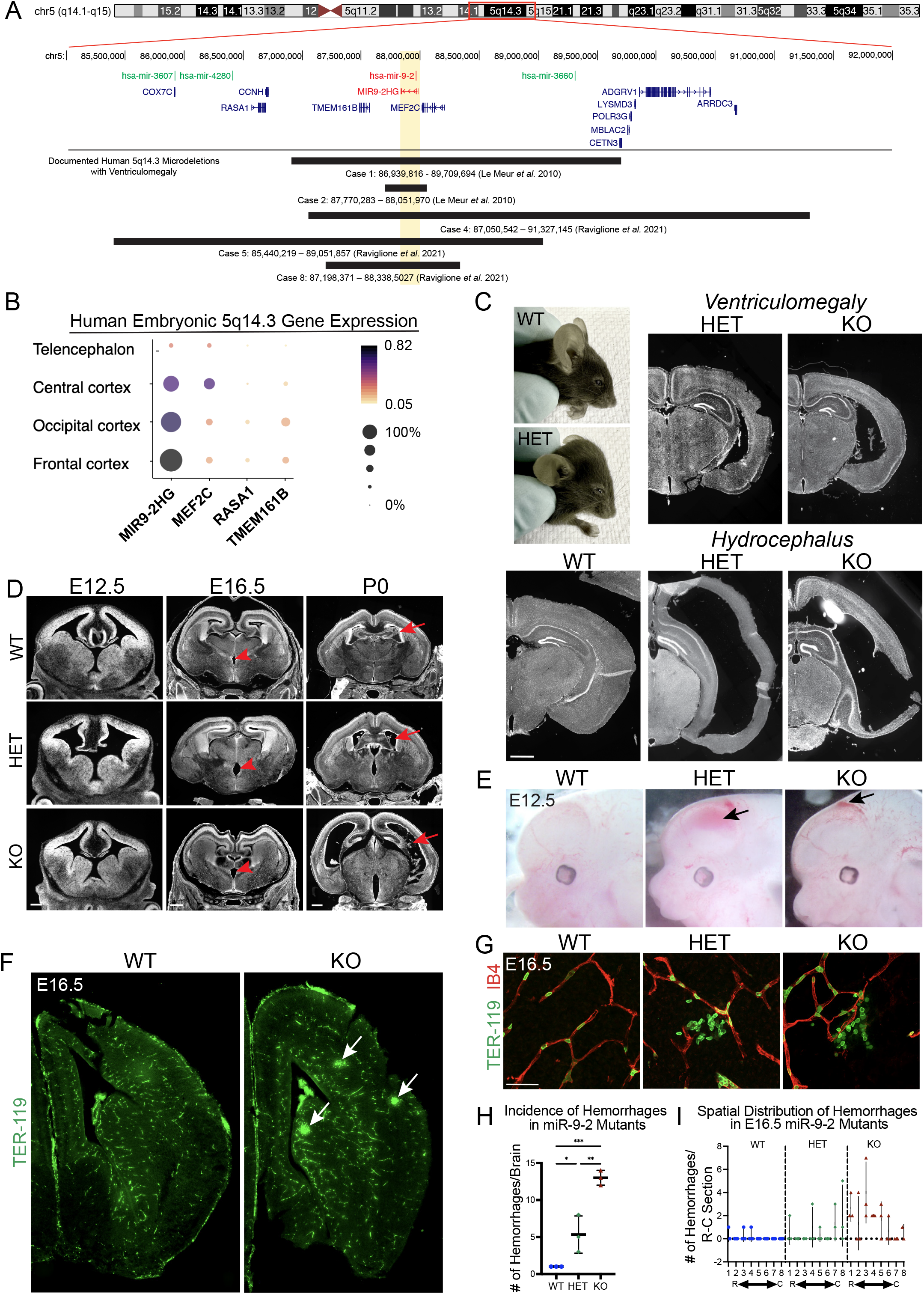
Loss of miR-9-2 causes ventriculomegaly/hydrocephalus with coincident cerebral hemorrhage in the developing mouse brain. **A**. UCSC Genome Browser track showing the human 5q14.3 locus containing *MIR-9-2*, its host gene *MIR9-2HG*, and documented human 5q14.3 microdeletions^6,8^. **B**. First trimester human embryonic single-cell RNA-seq gene expression data from Eze *et al*.^16^ showing relative expression of *MIR9-2HG* and 5q14.3 Microdeletion Syndrome associated genes. **C**. Whole head photographs of 3-week-old miR-9-2 WT and HET mice showing doming of cranium in HETs consistent with hydrocephalus. Accompanying DAPI-stained coronal brain sections of 3-week-old miR-9-2 HETs, KOs and WT corroborate a range of hydrocephalic phenotypes. Scale bar = 1mm. **D**. DAPI-stained coronal brain sections at E12.5, E16.5 and P0 show progression of ventriculomegaly phenotype, beginning with enlargement of third ventricle (red arrow heads) at E16.5 followed by enlargement of lateral ventricles (red arrows) by P0. Scale bars at: E12.5 = 250μm, E16.5 and P0 = 500μm. **E**. Whole head photographs showing that E12.5 miR-9-2 mutant embryos often present with intraventricular (HET) and subdural (KO) hemorrhages. The occurrence of either is not mutually exclusive by genotype. **F**. Immunohistology showing clusters of extravasated TER-119^+^ red blood cells, indicative of microhemorrhages within the E16.5 KO brain parenchyma. **G**. High-magnification confocal images showing extravasated TER-119^+^ red blood cells surrounding IB4-stained blood vessels in E16.5 miR-9-2 mutants. Scale bar = 50μm. **H**. Total hemorrhage counts among E16.5 miR-9-2 mutants and controls spanning rostral to caudal coronal brain sections. One-way ANOVA with Tukey’s multiple comparisons test revealed significant differences among all groups: P = *0.0337, **0.0023, ***0.0002. **I**. Numbers of hemorrhages by rostral to caudal (R-C) position among E16.5 miR-9-2 mutants and controls. Two-way ANOVA revealed no significant difference: P = 0.4179. (**H**,**I**: n = 3 animals per genotype, n = 3 slides per animal containing 6-8 coronal sections each).

To assess the potential role of *miR-9-2* (S.Fig. 1A) in brain development and disease, we generated germline *miR-9-2* knock-out (KO) mice (S.Fig. 1B). We observed that heterozygous (HET) mice develop enlarged skulls consistent with hydrocephalus, a common feature of 5q14.3 Microdeletion Syndrome, by approximately 3 weeks of age (Fig. 1C) at a frequency of 27% in males and 32% in females (S.Fig. 1C) showing that, similar to 5q14.3 Microdeletion Syndrome, *miR-9-2* is haploinsufficient in brain development. Although homozygous knockout (KO) mice are born alive and at expected Mendelian ratios, 98% of KO pups die within 1-2 days of birth (S.Fig. 1D). Those that do survive up to 3 weeks of age have a small body size (S.Fig. 1E). To determine if hydrocephalus develops embryonically, we compared *miR-9-2* wild-type (WT), HET, and KO littermates histologically from embryonic days (E) 12.5, E16.5 and postnatal day (P)0 (Fig. 1D) coincident with the timing of *miR-9-2* expression^3^. Embryonic brain development appears normal among *miR-9-2* mutants (HETs and KOs) at E12.5. However, by E16.5, enlargement of the third ventricle is frequently observed in HET and KO brains. By P0, enlargement of lateral and third ventricles is evident in 100% of *miR-9-2* mutants and is particularly severe in *miR-9-2* KOs (Fig. 1D).

In mouse models, congenital hydrocephalus is most commonly attributed to disturbed cerebrospinal fluid (CSF) circulation from disruption of ciliated ependymal cells lining the cerebral ventricles^17-19^ or blockage of flow through the cerebral aqueduct^18^. However, histological examination of *miR-9-2* mutant brains revealed no defects in these structures (S.Fig. 1F-H). Additionally, we see enlarged ventricles in HET and KO brains before ependymal cilia normally develop in the first postnatal week^20^, suggesting disrupted ependyma and aqueduct blockage are not initiating factors (Fig. 1D).

Notably, we did often observe cerebral hemorrhages in HET and KO brains prior to ventricular enlargement as early as E12.5 (Fig. 1E). Intraventricular hemorrhage (IVH) is a known cause of congenital hydrocephalus in human births^21^, we therefore assessed the frequency and distribution of hemorrhages in mutant brains at E12.5 and E16.5 using vascular (IB4) and red blood cell (TER-119) markers. At E12.5, the hemorrhages appeared to be intraventricular or subdural (Fig. 1E). However, by E16.5, we frequently observed deep, parenchymal microhemorrhages spanning brain regions (Fig. 1F-I), suggesting widespread and stochastic vascular destabilization in *miR-9-2* HETs and KOs that could contribute to the hydrocephalus phenotype observed in this mouse model.

To further define the cellular and molecular consequences of *miR-9-2* loss, we performed single-nucleus RNA-sequencing (snRNA-seq) on *miR-9-2* WT and KO dorsal forebrains (Fig. 2A, S.Fig. 2A,B) prior to and coincident with onset of enlargement of the cerebral ventricles (E12.5 and E16.5). We identified cell classes based on canonical marker expression (S.Fig. 2C,D) and confirmed recovery of cell classes in the expected proportions^22,23^ (Fig. 2B,C). The miR-9 family has been shown to regulate cell cycle progression and influence cell class proportions^2,3^. However, we found that loss of *miR-9-2* alone in the developing cortex did not cause significant changes to cell cycle (S.Fig. 3A,B) or cell class proportions (Fig. 2C, S.Fig. 3C,D) potentially due to partial redundancy with the other miR-9 family members.

**Fig. 2.**
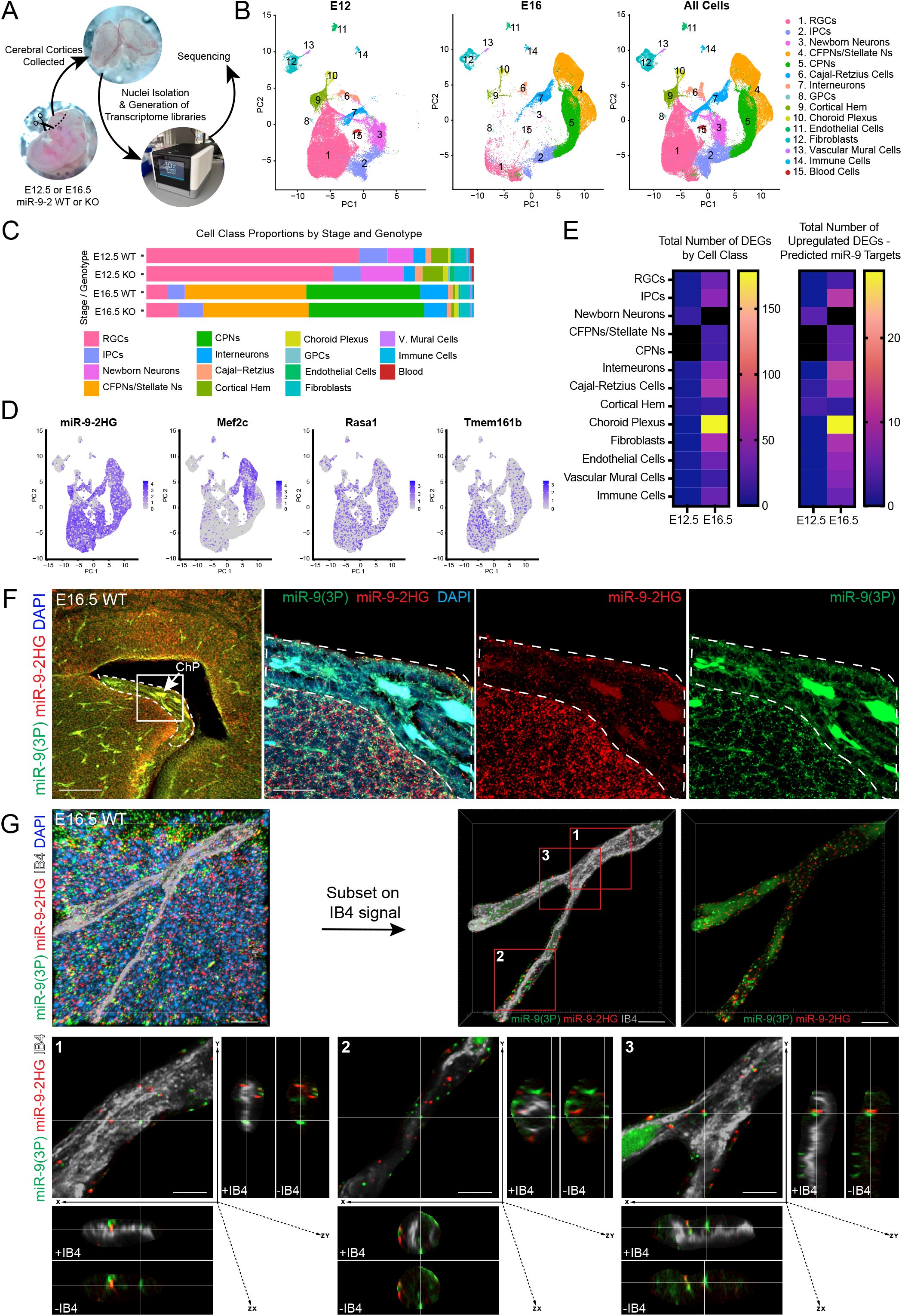
Single-nucleus RNA-seq of miR-9-2 KO and control cortices reveals transcriptional dysregulation among developing brain cell classes. **A**. Schematic showing snRNA-seq workflow. **B**. UMAP representations of harvested cortical cell types collected at E12.5 and E16.5 timepoints for integrated miR-9-2 WT and KO samples. **C**. Stacked bar plots showing captured cell class proportions between E12.5 and E16.5 miR-9-2 WT and KO samples. **D**. Gene expression feature plots showing relative expression of *miR-9-2HG, Mef2c, Rasa1* and *Tmem161b* among all E12.5 and E16.5 WT cell populations. **E**. Heat maps showing numbers of total differentially expressed genes and numbers of derepressed predicted miR-9-2 target genes per cell class per timepoint between miR-9-2 WT and KO littermates at FDR ≤ 0.05 and -0.5 ≤ log_2_FC ≥ 0.5. Black boxes indicate *no data*. **F-G**. *In situ* hybridization confocal images showing expression of both mature *miR-9(3’ strand)* and *miR-9-2HG* transcripts in E16.5 **(F)** choroid plexus and **(G)** IB4^+^ capillary-caliber cortical blood vessel. The blood vessel was skeletonized and isolated to clarify *miR-9* puncta expression within IB4^+^ signal. Subsetted images 1-3 show high-magnification examples of *miR-9(3p)/miR-9-2HG* puncta within IB4 signal along stained vessel. Scale bars (F) = 100 μm 1^st^ panel, 25μm 2^nd^ panel; (G) = 10μm large upper panels, 5μm subsetted panels 1, 2, 3. RGCs = radial glial cells, IPCs = intermediate progenitor cells, CFPNs = corticofugal projection neurons, CPNs = cortical projection neurons, GPCs = glial precursor cells, CH = cortical hem, ChP = choroid plexus.

We therefore hypothesized that loss of *miR-9-2* drives cerebral hemorrhages and enlarged ventricles by disrupting cell-type-specific transcriptional programs. We expected vascular endothelial cells, which form the developing vasculature, and choroid plexus cells, which secrete and regulate CSF, to be most strongly affected. Indeed, both these cell classes express *miR-9-2* (Fig. 2D,F,G) and show significantly affected gene expression in KO brains (Fig. 2E). However, we also observed significant dysregulation of transcriptional programs across many cell classes, suggesting that loss of *miR-9-2* has widespread consequences on the developing brain (Fig. 2E). Indeed, as in humans, murine *miR-9-2HG* is more highly and broadly expressed among neural cell types compared to other 5q14.3 Microdeletion Syndrome-implicated genes (Fig. 2D). In particular, we see significantly altered gene expression in neural progenitor and newborn neuron populations supporting newly proposed roles for these cell classes in the pathobiology of hydrocephalus^24^.

At E12.5, before the peak of *miR-9-2* expression in the brain, the transcriptional consequences of *miR-9-2* loss are subtle. We saw a de-repression of computationally predicted *miR-9-2* targets^25,26^ in early born cell types known to be important for patterning of the developing cortex, including Cajal-Retzius and Cortical Hem cells and newborn neurons (Fig. 3A-C). These included transcription factors like *Pbx3* in Cajal-Retzius Cells and newborn neurons, *Lmo3* in Cortical Hem cells, and *Zfhx4*, and *Foxp2* in newborn neurons (Fig. 3A-C) that could cause secondary dysregulation of neural gene networks. Only modest dysregulation of gene expression is observed among other cell classes at this timepoint (S.Fig. 4).

**Fig. 3.**
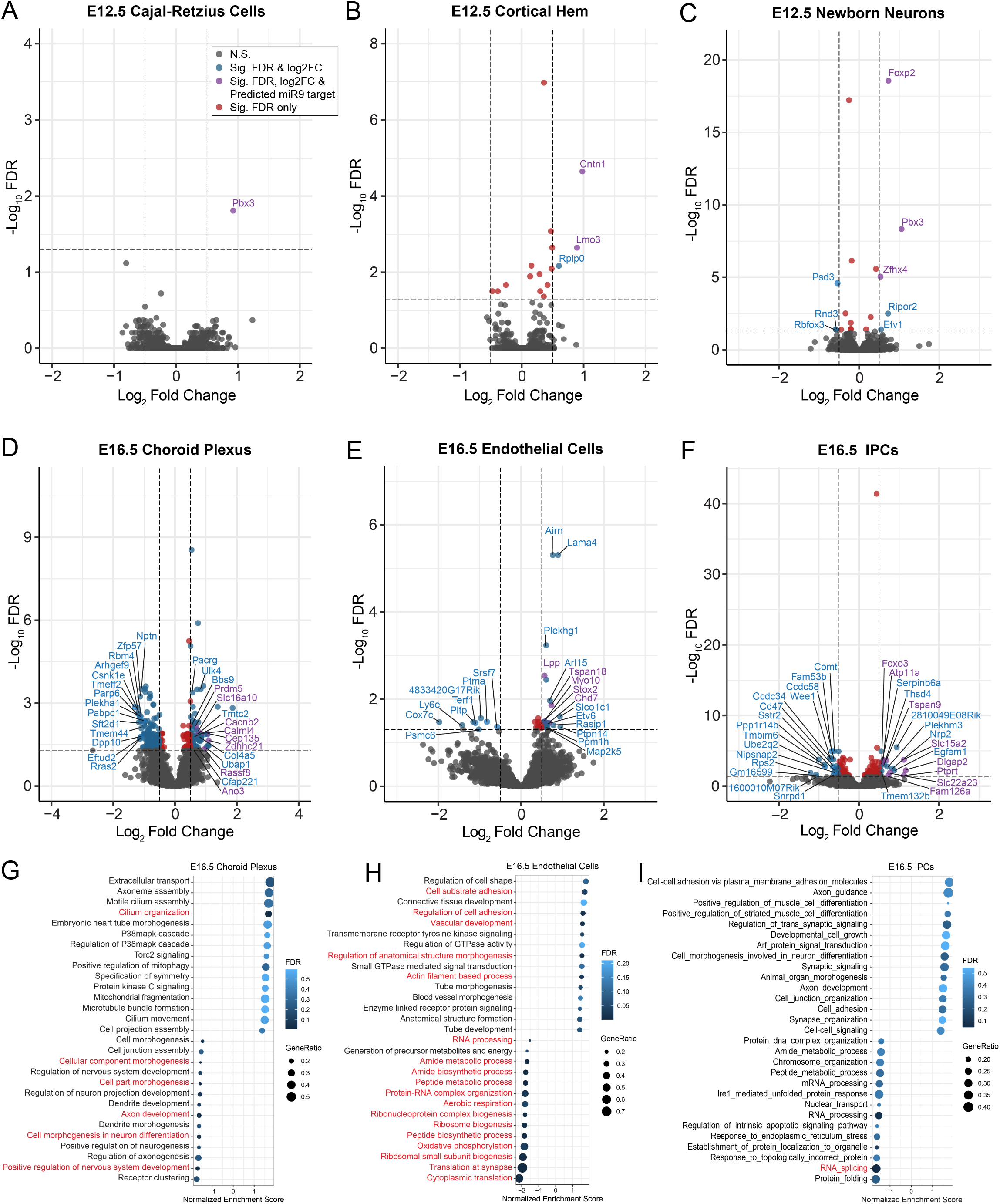
miR-9-2 KO cortices display DEGs across a wide range of cortical cell types. **A-F**. Volcano plots showing differentially expressed genes at FDR ≤ 0.05 and -0.5 ≤ log_2_FC ≥ 0.5 for (**A**) E12.5 Cajal-Retzius cells, (**B**) E12.5 cortical hem, (**C**) E12.5 newborn neurons, (**D**) E16.5 choroid plexus, (**E**) E16.5 vascular endothelial cells and (**F**) E16.5 intermediate progenitor cells. **G-I**. FGSEA results for enriched GO-BPs for (**G**) E16.5 choroid plexus, (**H**) E16.5 vascular endothelial cells and (**I**) E16.5 intermediate progenitors showing top and bottom 14 most differentially enriched BPs. BPs in red font indicate pathways significant at FDR ≤ 0.05 and p ≤ 0.05. All other pathways are significant at p ≤ 0.05. FGSEA = fast gene set enrichment analysis, GO-BP = gene ontology biological processes.

By E16.5, after the peak of *miR-9-2* expression, the consequences of *miR-9-2* loss were more dramatic and widespread (Fig. 3D-F, S.Fig. 5). Choroid plexus cells showed the highest number of differentially expressed genes and the highest number of derepressed *miR-9-2* targets (Fig. 2E, 3D) consistent with the hydrocephalus phenotype. Vascular endothelial cells also have significant changes in gene expression and derepressed *miR-9-2* targets, consistent with cerebral hemorrhages (Fig. 2E, 3E). We further sub-clustered choroid plexus and vascular endothelial cells to more closely assess *miR-9-2HG* expression and found that within each class *miR-9-2* expression was enriched in a distinct subset of cells (S.Fig. 6). In choroid plexus, this subset appeared to be epithelial cells based on *Ttr* expression as well as β-gal expression from the original *miR-9-2* knock-in allele^27^ (S.Fig. 6A). In vascular endothelial cells, this subset appears to have a capillary-venule identity, consistent with our observation that hemorrhages are typically associated with small vessels at this stage (S.Fig. 6B,C). Other cell classes that showed significant transcriptional changes included developing neural cells (Fig. 3F, S.Fig. 5), further supporting a role for progenitor and precursor cells in congenital hydrocephalus^24,28-30^, and non-neural cells such as perivascular fibroblasts and immune cells that may also contribute^31^.

Given the widespread effects of *miR-9-2* loss, it is beyond the scope of this study to determine which affected transcripts in each cell class are drivers of specific phenotypes. However, to understand overall changes to cell biological processes, we performed Gene Set Enrichment Analysis for each affected cell class. Affected genes in choroid plexus at E16.5 were associated with *extracellular transport*, potentially related to CSF secretion, and *cilia development*, which could reflect disrupted embryonic signaling or motility in newly described choroid plexus cilia^32^ (Fig. 3G). In vascular endothelial cells, gene expression changes were associated with *cell adhesion* and *vascular development* (Fig. 3H). To directly assess cerebral vascular integrity, we perfused E16.5 embryos with a fluorescent vascular tracer as previously described^33^ and indeed detected sites of active hemorrhage (S.Fig. 7A), which is supported by prior studies of miR-9 family in neurovascular development^34^. However, we did not detect overall changes in vascular network organization (S.Fig. 7B-F). Among remaining cell classes, the associated biological processes were highly diverse suggesting a role for *miR-9-2* in the maintenance of cell-type-specific transcriptional programs and suppression of non-neural pathways (Fig. 3I, S.Fig. 8).

In conclusion, the results of this initial study reveal critical roles for *miR-9-2* that have immediate relevance for understanding neurodevelopmental disorders. This work extends beyond previous studies by showing that, among other miR-9 family members, *miR-9-2* is a unique, non-redundant regulator of cell-type-specific gene expression and is essential for the proper development of cerebral vasculature, choroid plexus, and cortical neurons. Interestingly, *miR-9-2* mouse mutants display phenotypes similar to those of 5q14.3 Microdeletion Syndrome, a human neurodevelopmental condition that commonly arises from a loss of *MIR-9-2* and other nearby genes, but that has not been previously connected to *MIR-9-2* function. Furthermore, we show that proper *miR-9-2* dosage is required, as 100% of HET and KO animals have ventriculomegaly, which may help explain genetic risk for autism in a cortical enhancer of *MIR-9-2* expression^4,5^. Lastly, we find that loss of *miR-9-2* leads to three major risk factors for congenital hydrocephalus: cerebral hemorrhages, disrupted choroid plexus, and dysregulated neural cell development. Thus, *MIR-9-2* warrants closer attention as a potential contributor to human neurodevelopmental disorders and possibly as a therapeutic target for these intractable conditions.

## Acknowledgements

We would like to thank the animal care team in the Office of Animal Care at Seattle Children’s Research Institute (SCRI). We also thank Drs. Stephanie Bonney and Andy Shih for their consultations and feedback throughout this study. S.P.F. was supported by a fellowship from the NIH/NINDS (F32NS127946). M.A. was supported by fellowships from SCRI Summer Scholars Program, the Undergraduate Summer Fellows Program from the Institute for Stem Cells and Regenerative Medicine at the University of Washington (UW) and the Mary Gates Research Scholarship from UW. K.L.C. was supported by a fellowship from the NIH/NEI (F31EY035932). D.M. was supported by PHS NRSA T32GM007270 from NIGMS. T.J.C. and the Cherry lab are supported by funding from the Hydrocephalus Association Network for Discovery Science Innovator Award, the Lowy Medical Research Institute, and NIH grants (R01EY028584 and R01EY033364). We thank the Shih lab for sharing their vascular antibodies and stains and we thank Drs. Kimberly Aldinger, Lucas Cheadle, Jennifer Franks, Allison Paquette and Mr. Brendan McShane for their constructive feedback on this manuscript.

## METHODS

### Animals

Mice were housed in a specific pathogen-free facility approved by the AAALAC. All work with animals was compliant with Seattle Children’s Research Institutes IACUC protocols. MiR-9-2 KO mice were generated by crossing the *Mir9-2*^*tm1Mtm*^ conditional KO mice (RRID:MMRRC_036062-JAX)^27^ first to Rosa-FLP (RRID:IMSR_JAX:009086)^35^ then to E2a-Cre (RRID:IMSR_JAX:003724) ^36^mouse lines to create miR-9-2 heterozygous animals. Subsequently, miR-9-2 HET animals were back-crossed to C57BL/6 (Jackson Labs) animals for 10 generations to homogenize the strain background. The uniformity of the C57BL/6J genetic background was validated using the MiniMUGA genotyping array (NeoGen). MiR-9-2 HET x HET crosses were then performed to generate WT, HET and KO littermate cohorts for experiments and analyses. Timed pregnant breeding was utilized to generate embryonic samples with day of copulation plug noted as E0.5. Equal numbers of male and female mice were used for all analyses, with the exception of E16.5 WT female samples, of which one was discarded due to poor quality, reducing E16.5 WT representation to one female and two males (S.Fig. 2A).

### Immunohistochemistry and Imaging

All tissue for immunohistochemistry was collected and fixed in cold 4% paraformaldehyde overnight at 4°C and then cryoprotected in 30% sucrose in sterile 1X phosphate buffered saline (1X PBS) at 4°C for 2-4 days. Tissue was then embedded and frozen in TissueTek O.C.T. (Sakura #4583) and sectioned on a Leica CM 3050 S cryostat (40-50μm sections) and mounted on adhesive glass microscope slides. Prior to staining, slides were incubated in a 60ºC oven for up to 3 hours to enhance tissue adhesion. Slides were washed 3x in 1X PBS before and after incubation with primary and secondary antibodies. Prior to primary antibody incubation, sections were subjected to sodium citrate (0.01M Na-Citrate, pH 6.0) antigen retrieval in a pressure cooker for 8-10 minutes on the “high” setting. All antibodies were prepared in 1X PBS containing 5% bovine serum albumin (BSA) and 0.2% Triton X-100. DAPI was used to counterstain nuclei.

The following antibodies and stains were used as follows: Rabbit anti-ZO-1 (1:250; Abcam Cat# ab221547, RRID:AB_2892660); Mouse anti-Acetylated Tubulin (1:250; Sigma-Aldrich Cat# T6793, RRID:AB_477585); Goat anti-Sox9 (1:250; R and D Systems Cat# AF3075, RRID:AB_2194160); Rat anti-TER-119 (1:200; Thermo Fisher Cat# 14-5921-82, RRID:AB_467727); Rabbit anti-B-galactosidase (1:200; Sigma-Aldrich Cat# AB1211, RRID:AB_177320); Donkey anti-Rabbit Alexa Fluor 555 (1:500; Thermo Fisher Cat# A-31572, RRID:AB_162543); Donkey anti-Mouse Alexa Fluor 555 (1:500; Thermo Fisher Cat# A-31570, RRID:AB_2536180); Donkey anti-Goat Alexa Fluor 488 (1:500; Thermo Fisher Cat# A-11055, RRID:AB_2534102); Donkey anti-Rat Alexa Fluor 488 (1:500; Thermo Fisher Cat# A-21208, RRID:AB_141709); Isolectin GS-IB_4_ -Alexa Fluor 647 conjugated (1:50; Thermo Fisher Cat# I32450).

All microscopic imaging was performed using either an Olympus APX100 fluorescence microscope or Zeiss LSM 900 or 980 confocal microscopes.

### Image Analysis

Cerebral hemorrhage counts were performed on E12.5 and E16.5 brain sections: n = 3 mice per genotype, n = 3 slides per animal containing 6-8 coronal sections each representing the full rostral-to-caudal axis. Absolute numbers of hemorrhages were counted in whole section slide scanner images using ImageJ/FIJI.

Vascular network analysis was performed using AngioTool^37^ on isolectin-B4-stained E12.5 and E16.5 brain sections. Images were first cropped to cerebral cortex regions of interest (i.e. medial and lateral cortex) using ImageJ/FIJI then imported into AngioTool. IB4 labeled blood vessel images were then scaled, skeletonized, and measured for vessel length, vascular density, and junction density.

### 10X Genomics Transcriptome Library Generation

Cerebral cortices were harvested from E12.5 and E16.5 embryos (n = 2 WT or KO/sex; total n = 4/genotype) in ice cold sterile filtered 1X PBS and immediately flash frozen in liquid nitrogen in nuclease-free microcentrifuge tubes then stored in vapor phase liquid nitrogen until processing for snRNA-seq.

For nuclei isolation, samples were thawed on ice and processed via a modified 10X Genomics Nuclei Isolation from Mouse Brain Tissue protocol (CG000212 Rev. B). 500 µl of cold 0.1X Lysis Buffer (10 mM Tris-HCl pH 7.4, 10 mM NaCl, 3 mM MgCl2, 0.01% Tween-20, 0.01% IGEPAL CA-360, 1% BSA, 1 U/µl RNase Inhibitor (Roche #03335402001)) was added to each sample-containing tube and triturated 15-20 times. The suspensions were further homogenized using dounce homogenizers by applying 10 dounces each with an A then B pestle. Following transfer of each sample to a new tube, the samples were incubated on ice for 5 minutes. Sample suspensions were pipette-mixed 10 times, then incubated on ice for 10 minutes. 500 µl of cold Wash Buffer (10 mM Tris-HCl pH 7.4, 10 mM NaCl, 3 mM MgCl2, 1% BSA, 0.1% Tween-20, 1 U/µl RNase Inhibitor) was then added, and samples were pipette-mixed 5 times and then passed through a 40 µm Flowmi Cell Strainer into a fresh tube. Dissociated sample nuclei were then washed 2 times by centrifugation pelleting at 500 rcf in a 4ºC swinging-bucket centrifuge and resuspension in 750 µl cold Wash Buffer. Sample nuclei concentrations were determined using propidium iodide and a Countess II FL Automated Cell Counter. Nuclei were then centrifuged at 500 rcf for 5 minutes at 4ºC. The supernatant was discarded and the nuclei were resuspended in cold Nuclei Buffer (1X 10X Genomics Nuclei Buffer) to achieve a nuclei concentration range of 700-1200 nuclei/ µl, which is recommended for targeting 10,000 cells in the 10x Genomics snRNA-seq protocol. The final nuclei resuspension concentration was confirmed using the Countess II, followed by completion of the 10X Genomics Chromium Next GEM Single Cell 3’ v3.1 snRNA-seq protocol.

### Sequencing

All sample transcriptome libraries were sequenced by the Northwest Genomics Center at the University of Washington. Samples were sequenced per 10X Genomics’ recommended cycles and sequencing depth using either a NovaSeq SP or NovaSeq S1 100-cycle flow cell, according to the number of samples submitted for sequencing (SP for 4 or S1 for 5-9 samples). Data were formatted and returned for analysis as FASTQ files.

### snRNA-seq and Statistical Analyses

All data processing and statistical analysis was performed using R^38^. Sequencing reads were aligned to the GRCm39/mm39 and UMI counts per gene and cell were quantified using Cellranger 7.2.0 (10X Genomics). Cellbender 0.3.0^39^ was used to remove background UMI counts and DoubletFinder 2.0.6^40^ was used to remove doublets. Seurat v5^41^ was then used for quality control, filtering, normalization, ordination, integration, and cell cycle scoring. Cells were retained for analysis if they had between 1000 and 15,000 UMIs, at least 200 genes detected, and less than 5% of UMIs were mitochondrial, resulting in 84,128 cells. Genes were retained for analysis if they were detected in at least 20 cells. Seurat’s ‘NormalizeData’ function was used for data normalization, and the 2000 most variable genes were used for principal components analysis. Based on preliminary clustering, integration was run across four batches of libraries, each containing equal representation of sex and genotype (1 WT: 1 KO each of male and female samples) per stage, using the RPCA method, which generated a set of reduced dimensions used for cell clustering and Uniform Manifold Approximation and Projection (UMAP). Samples were distributed by sex, stage, and timepoint across five batches, and integration was run across batches of libraries using the Reciprocal PCA method from the Seurat package, after which cells were re-clustered. Cell clusters were labeled and manually grouped using marker gene expression. Nebula^42^ was then used for differential gene expression analysis within cell type and stage for genes that had a non-zero count in at least 10% of cells, comparing miR-9-2 WT to KO cells using sample as a random effect with a Benjamini-Hochberg corrected p-value threshold of 0.05. Volcano plots were generated using the Enhanced Volcano^43^(version 1.4.0) package to show differentially expressed genes per cell class and timepoint. The list of input genes and their associated log fold changes and false discovery rates (FDR) was generated from Nebula^42^. The significance threshold was set at -log10 FDR ≤ 0.05, and the significant log_2_ fold change was set at ±0.5. Predicted miR-9-2 target genes were identified using TargetScan^25,26^. The FGSEA R package^44^ was used to test for enrichment of gene sets within the Gene Ontology (GO) biological processes gene sets. To rank genes for FGSEA within cell type and stage, the log-fold-change was multiplied by the negative log base 10 of the P-value^45^. Differential cell type abundance at each stage, between wild type and knockout cells, was determined by performing a quasi-likelihood F-test and fitting a negative binomial generalized linear model to the cell count data using EdgeR 4.4.2^46^ and using a Benjamini-Hochberg corrected p-value threshold of 0.05. Cluster subsetting was done by isolating the cluster of interest and using Seurat for ordination, integration, PCA, clustering, and UMAP, as described above.

### Embryonic Cerebral Hemorrhage Detection via Fluorescent Dextran Perfusion

Control and mutant embryos (E16.5) from timed-pregnant dams were perfused with fluorescent tracer dextran as previously described^33^. Briefly, E16.5 embryos were surgically exposed from their anesthetized mother’s uterus, leaving the umbilical cord intact to continually supply blood to the embryos during the perfusion. Embryos were then injected with 5μl of lysine-fixable Texas Red dextran (70kD) (Thermo Fisher Cat# D1864) with a 300μl insulin syringe via their highly fenestrated livers and allowed to circulate through each embryo for 3 min. Embryo umbilical cords were then severed and embryo heads were drop fixed in cold 4% paraformaldehyde overnight at 4°C then processed for histological analysis as described above.

### Quantifications, Statistics and Plots

Statistical analyses for image quantifications were performed using GraphPad Prism 10.1.1. Comparisons of absolute values between groups of three were performed using one-way ANOVA with Tukey’s multiple comparisons test. Comparisons of trends between groups of three were performed using a two-way ANOVA. Statistical significance was set at adjusted p ≤ 0.05.

## FIGURE LEGENDS

**SFig. 1. Generation of the germline miR-9-2 KO mouse allele and phenotyping. A**. UCSC Genome Browser track showing the homologous mouse 5q14.3 locus (13qC3). **B**. Schematic showing the breeding strategy for germline deletion of miR-9-2. **C**. Survival rates among miR-9-2 HET male (73%) and female (68%) animals that do not develop severe hydrocephalus. Data comprises 511 animals from 166 litters. **D**. miR-9-2 WT, HET and KO birth and survival rates. **E**. Surviving miR-9-2 KOs display a reduction in overall body size compared to WT littermates. **F**. Immunohistology of E16.5 miR-9-2 WT, HET and KO mouse brains showing intact ZO-1^+^ ependyma along lateral, 3^rd^ and 4^th^ ventricles. Scale bar = 100μm. **G**. DAPI-stained P0 miR-9-2 WT, HET and KO mouse brains showing no evidence of aqueductal stenosis along the cerebral aqueduct. Scale bar = 100μm. **H**. Immunohistology of P7 miR-9-2 WT and HET mouse brains showing normal-appearing Acetylated Tubulin^+^ motile cilia on Sox9^+^ ependymal cells. Scale bar = 20μm.

**SFig. 2. Quality control metrics and cell cluster identification for miR-9-2 WT and KO snRNA-seq samples. A**. Cell count summaries for E12.5 and E16.5 miR-9-2 WT and KO snRNA-seq samples. **B**. Quality control metrics for all snRNA-seq samples, including numbers of genes per cell, numbers of transcripts per cell and percent mitochondrial DNA per cell. **C**. Gene expression dot plot showing cell class markers that define captured cortical cell classes for E12.5 and E16.5 miR-9-2 WT and KO samples. **D**. Gene expression feature plots showing spatial distribution of key cell class marker genes on combined E12.5 and E16.5 UMAPs. RGCs = radial glial cells, IPCs = intermediate progenitor cells, CFPNs = corticofugal projection neurons, CPNs = cortical projection neurons, GPCs = glial precursor cells.

**SFig. 3. Cell cycle and cell class proportions for miR-9-2 WT and KO snRNA-seq samples. A**,**C**. Stacked bar plots showing distribution of (**A**) cycling cells by mitotic phase for E12.5 and E16.5 miR-9-2 WT and KO RGCs and IPCs and (**C**) cell class proportions for all snRNA-seq samples. **B**,**D**. Differential abundance statistical analyses results for (**B**) RGC and IPC cell cycle phase abundance and (**D**) cell class abundance by sample. RGCs = radial glial cells, IPCs = intermediate progenitor cells.

**SFig. 4. Volcano plots showing DEGs for E12.5 neural cell types. A-C**. Volcano plots showing differentially expressed genes at FDR ≤ 0.05 and -0.5 ≤ log_2_FC ≥ 0.5 for (**A**) E12.5 RGCs, (**B**) E12.5 IPCs, and (**C**) E12.5 Interneurons. RGCs = radial glial cells, IPCs = intermediate progenitor cells.

**SFig. 5. Volcano plots showing DEGs for E16.5 neural cell types. A-H**. Volcano plots showing differentially expressed genes at FDR ≤ 0.05 and -0.5 ≤ log_2_FC ≥ 0.5 for (**A**) RGCs, (**B**) CFPNs/stellate neurons, (**C**) CPNs, (**D**) Cajal-Retzius cells, (**E**) interneurons, (**F**) cortical hem cells, (**G**) Fibroblasts, and (**H**) Immune cells. RGCs = radial glial cells, IPCs = intermediate progenitor cells, CFPNs = corticofugal projection neurons, CPNs = cortical projection neurons.

**SFig. 6. Subsets of choroid plexus and vascular endothelial cells express *miR-9-2*. A-B**. UMAPs of subclustered (**A**) choroid plexus and (**B**) vascular endothelial cells from snRNA-seq datasets showing *miR-9-2HG* expression in each cell type. Each cell cluster is shown by stage, subcluster and cell type-specific marker gene expression. (**A**) also shows choroid plexus miR-9-2HG-driven β-galactosidase expression in non-recombined miR-9-2 KO mice at P0. **C**. Gene expression dot plot from snRNA-seq endothelial cell cluster showing expression of known vascular zonation (Capillary-Arteriole, Capillary, or Capillary-Venule) marker genes among brain endothelial cells. Marker genes curated from Chen *et al*. ^47^. ChP: choroid plexus.

**SFig. 7. Comprised vascular integrity underlies seemingly normal patterning in miR-9-2 mutant embryo brains. A**. Confocal images showing leakage of 70kD fluorescent dextran coincident with extravasated TER-119^+^ red blood cells around IB4-stained blood vessels in E16.5 miR-9-2 HETs. Scale bar = 50μm. **B-C**. Confocal images showing IB4-stained blood vessels in DAPI-counterstained cerebral cortices at (**B**) E12.5 and (**C**) E16.5 for WT, HET and KO miR-9-2 embryos. Scale bar = 200μm. **D-F**. AngioTool Metrics showing quantifications for (**D**) average vessel length, (**E**) vascular density and (**F**) junctional density for IB4-stained blood vessel networks in E12.5 and E16.5 miR-9-2 mutant cortices. Quantifications comprised n = 4-5 animals and n = 2-4 images per animal spanning rostral-caudal axis of cerebral cortex. One-way ANOVA with Tukey’s multiple corrections tests revealed no significant differences between cohorts.

**SFig. 8. Differentially enriched biological pathways for E16.5 miR-9-2 KO cortical cell classes. A-E**. FGSEA results showing differentially enriched GO-BPs for E16.5 miR-9-2 KO neural cell classes, (**A**) RGCs, (**B**) CFPN/Stellate Neurons, (**C**) CPNs, (**D**) Cajal-Retzius cells, and (**E**) interneurons. Pathways in red font are significant at FDR ≤ 0.05 and p ≤ 0.05. All other pathways shown are significant at p ≤ 0.05. FGSEA = fast gene set enrichment analysis, GO-BP = gene ontology biological processes, RGCs = radial glial cells, CFPNs = corticofugal projection neurons, CPNs = cortical projection neurons

