## Supplementary figures and images for "*Loss of miR-9-2* Causes Cerebral Hemorrhage and Hydrocephalus by Widespread Disruption of Cell-Type-Specific Neurodevelopmental Gene Networks"

### Supplemental Figure 1

#### Supplemental Figure 1

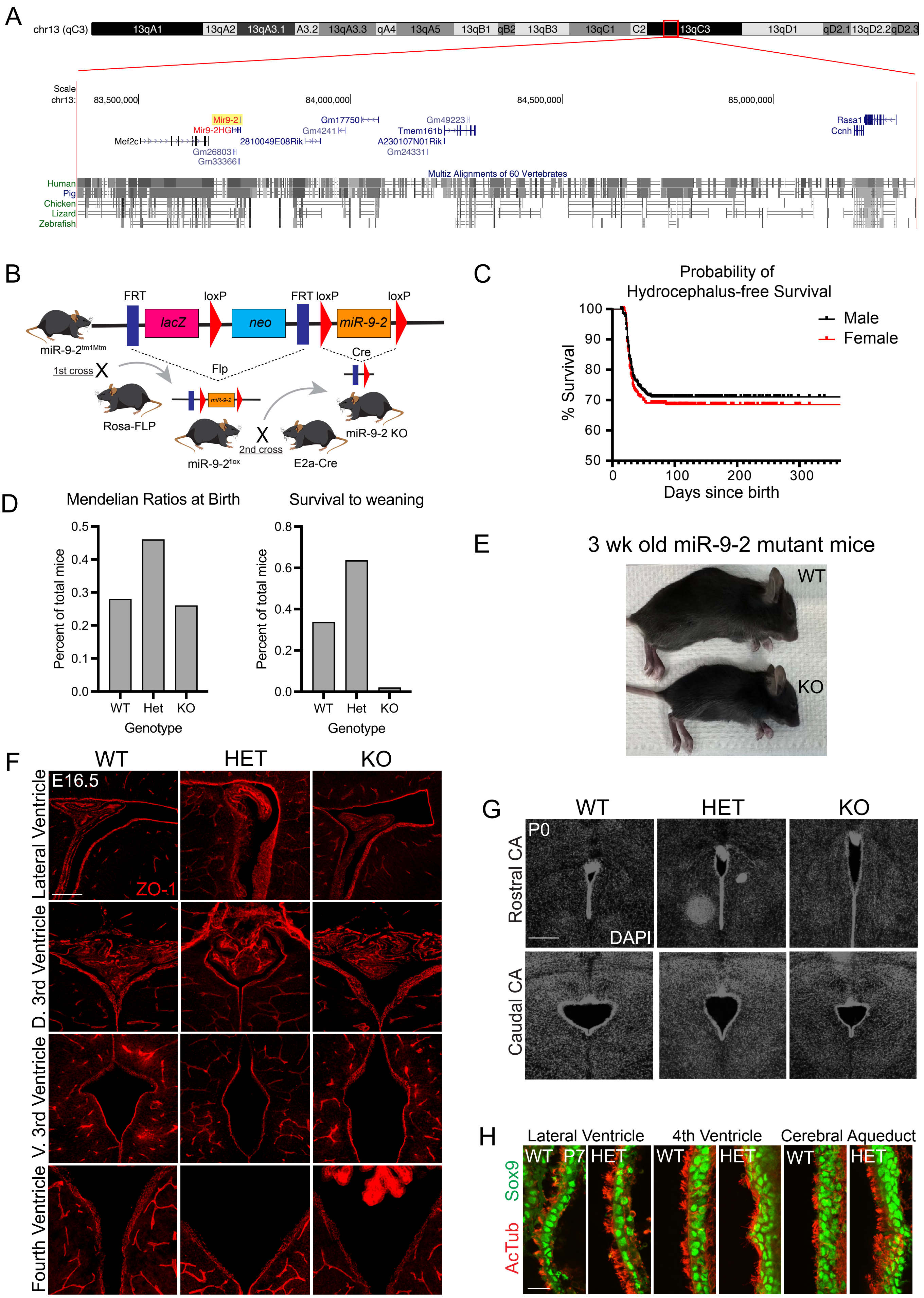

### Supplemental Figure 4

Supplemental Figure 4

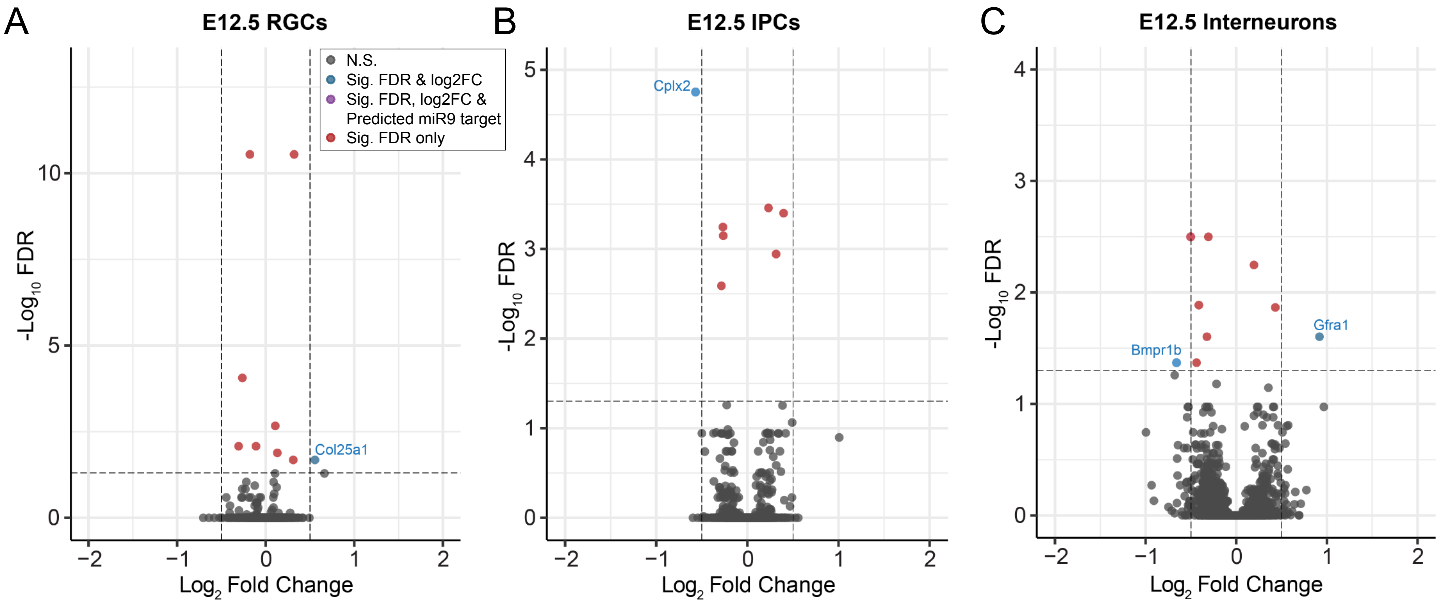

### Supplemental Figure 5

#### Supplemental Figure 5

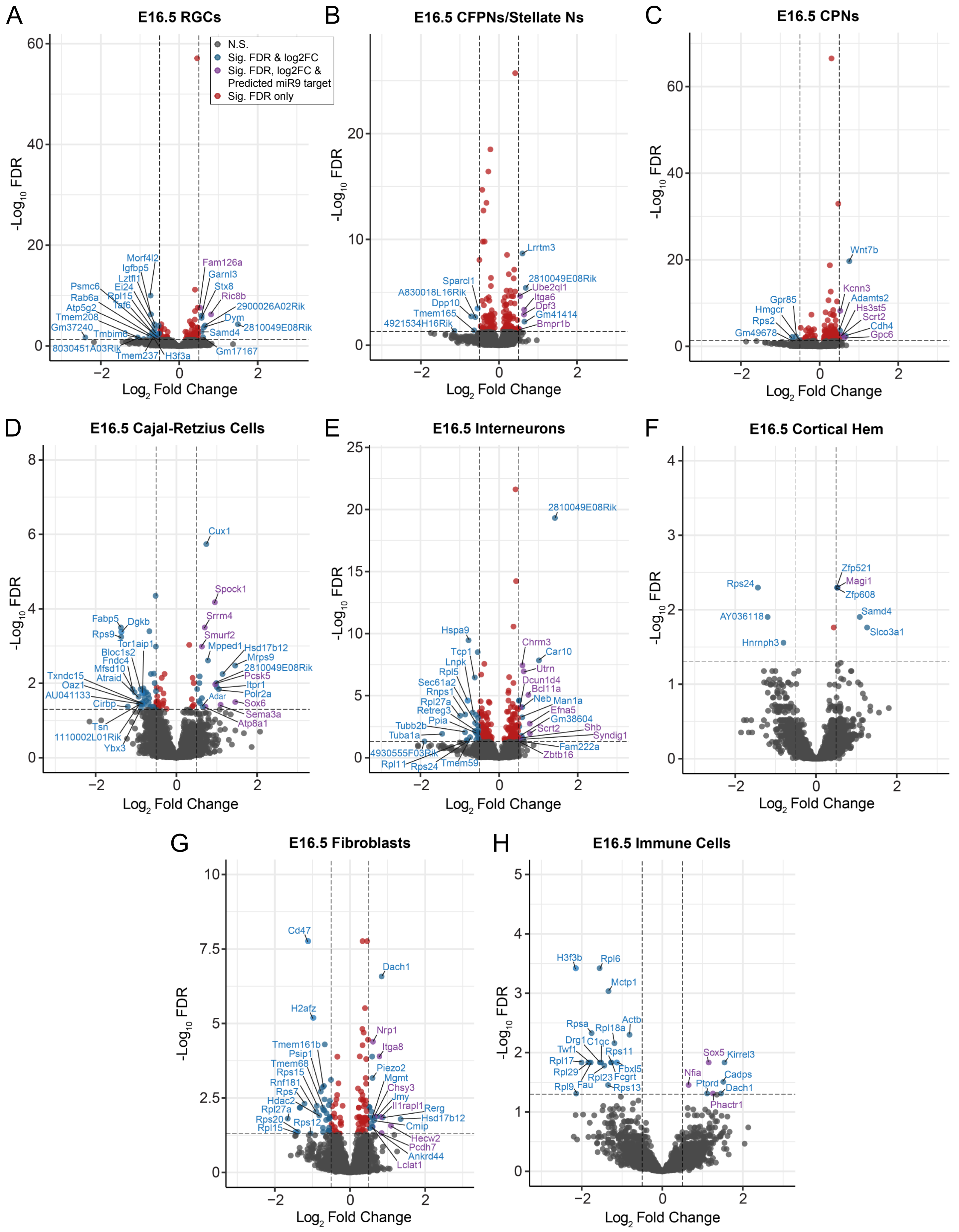

### Supplemental Figure 6

Supplemental Figure 6

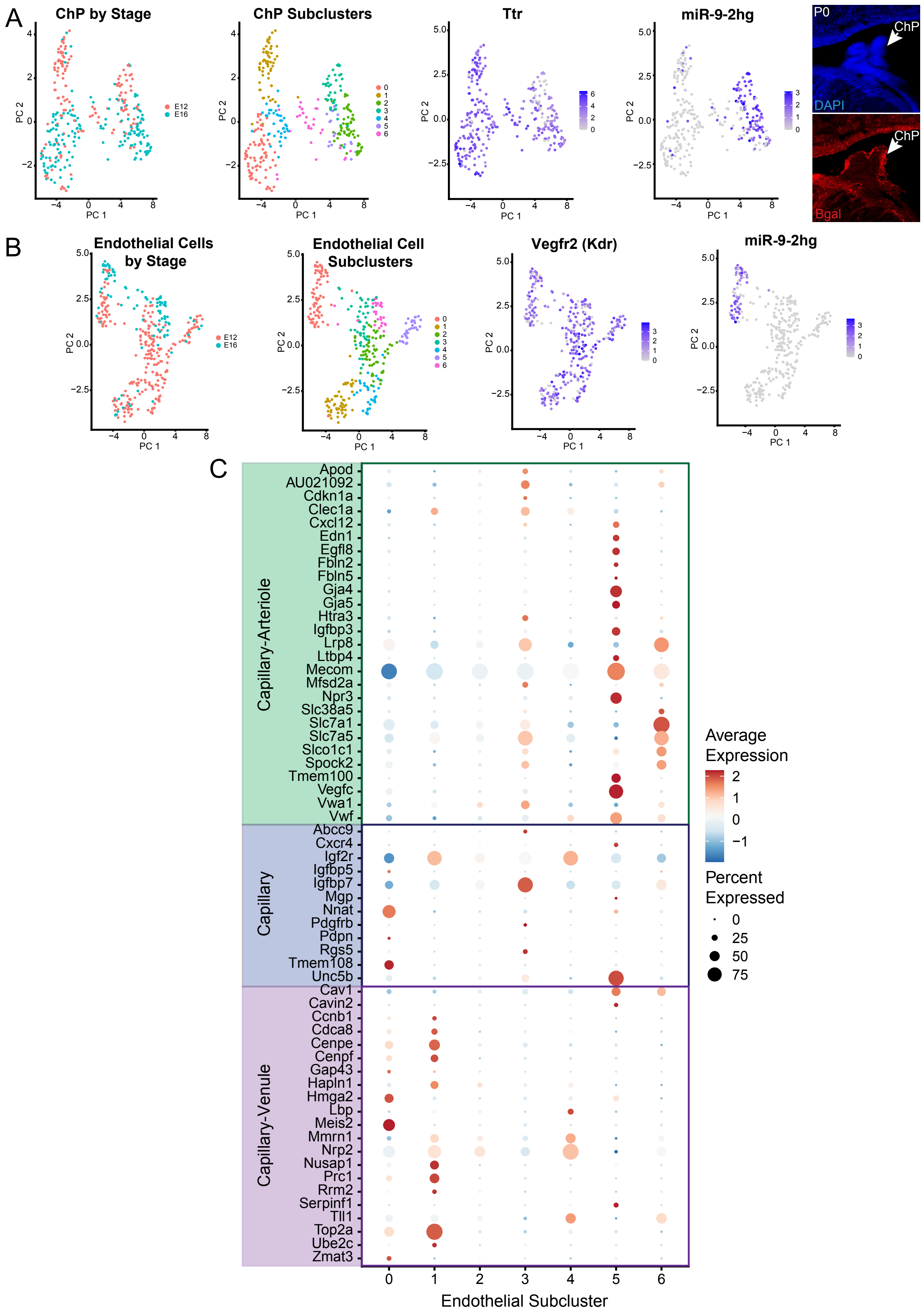

### Supplemental Figure 7

Supplemental Figure 7

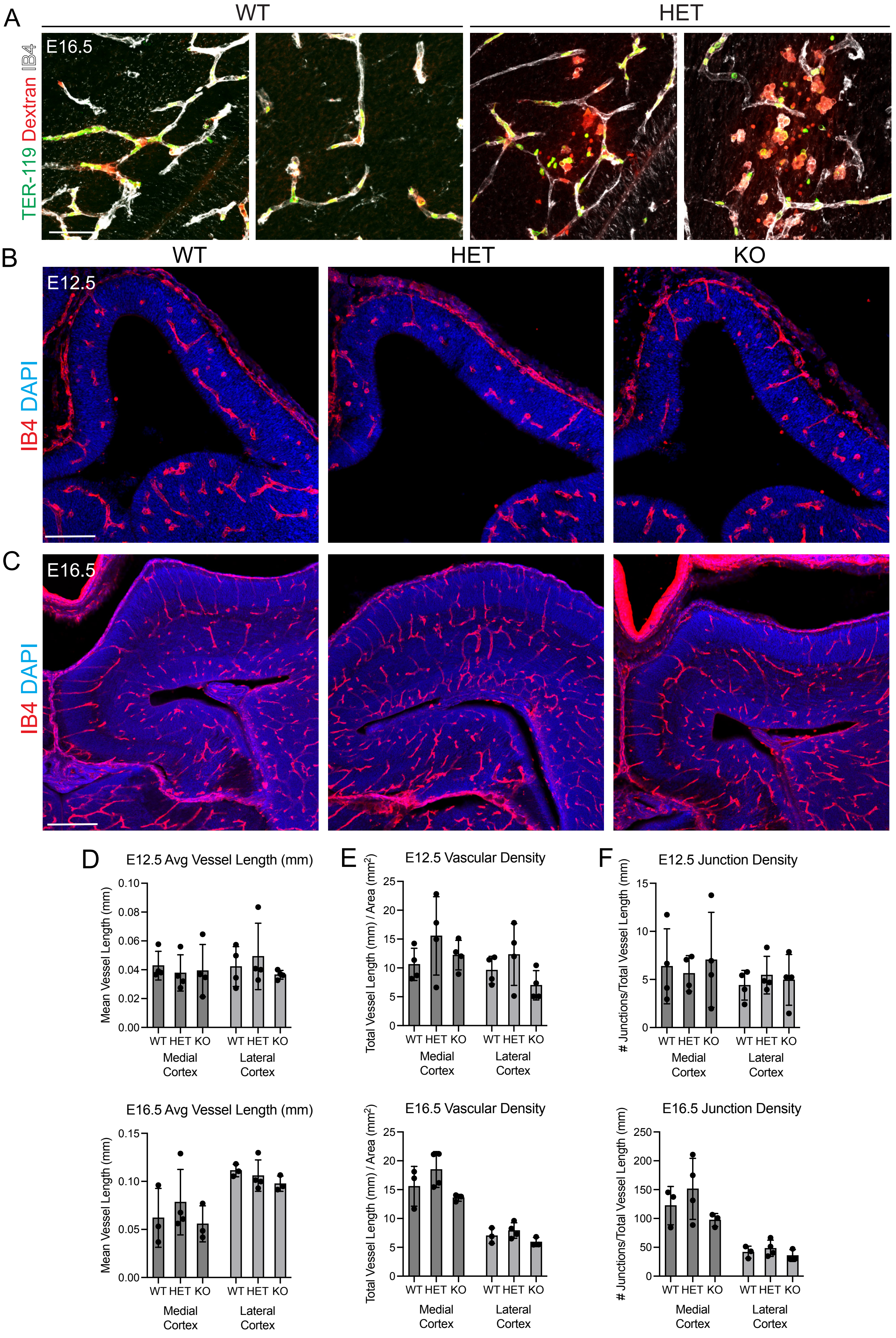

### Supplemental Figure 8

#### Supplemental Figure 8

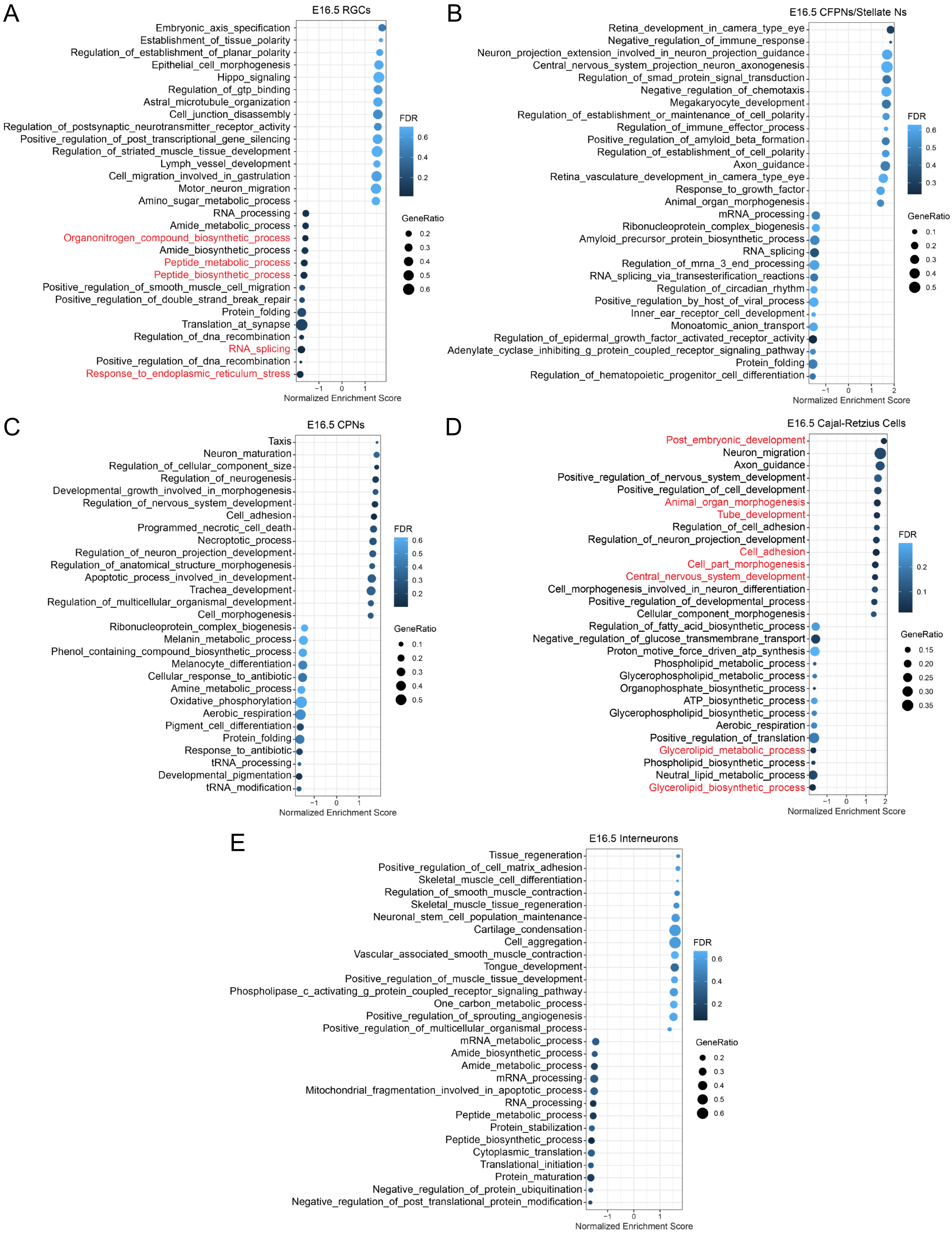
