## Supplemental Figure 2 for "*Loss of miR-9-2* Causes Cerebral Hemorrhage and Hydrocephalus by Widespread Disruption of Cell-Type-Specific Neurodevelopmental Gene Networks"

A

| Stage | Genotype | Sex | # Cells | Stage | Genotype | Sex | # Cells |
|---|---|---|---|---|---|---|---|
| E12.5 | WT | F1 | 6256 | E16.5 | WT | F2 | 5112 |
| | | F2 | 3844 | | | M1 | 6019 |
| | | M1 | 5488 | | | M2 | 6182 |
| | | M2 | 5337 | | | | |
| | KO | F1 | 7424 | | KO | F1 | 7862 |
| | | F2 | 4685 | | | F2 | 4222 |
| | | M1 | 3896 | | | M1 | 6951 |
| | | M2 | 6288 | | | M2 | 4562 |
| Total | | | | | | | 84,128 |

B

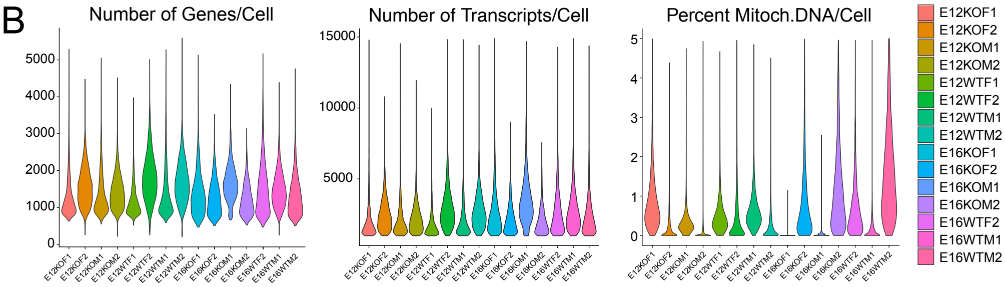


C

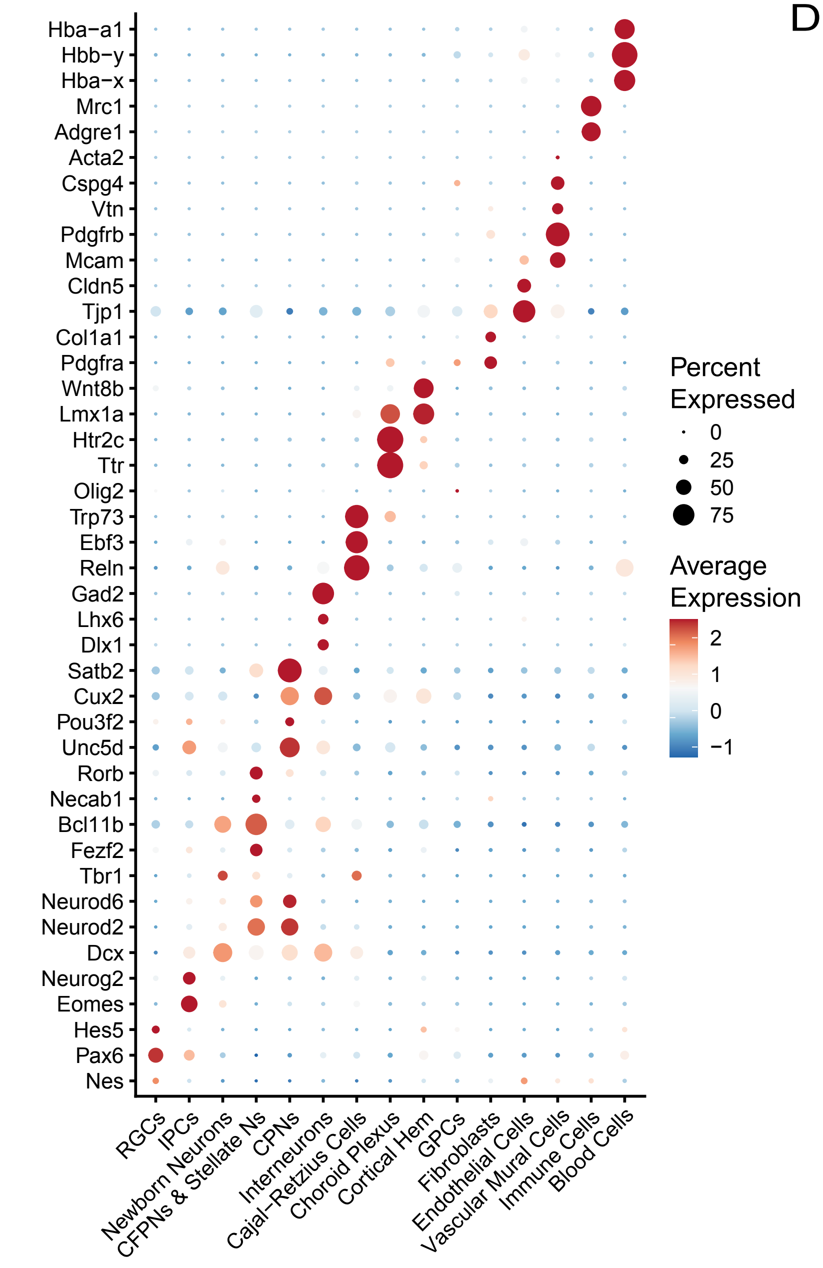


D

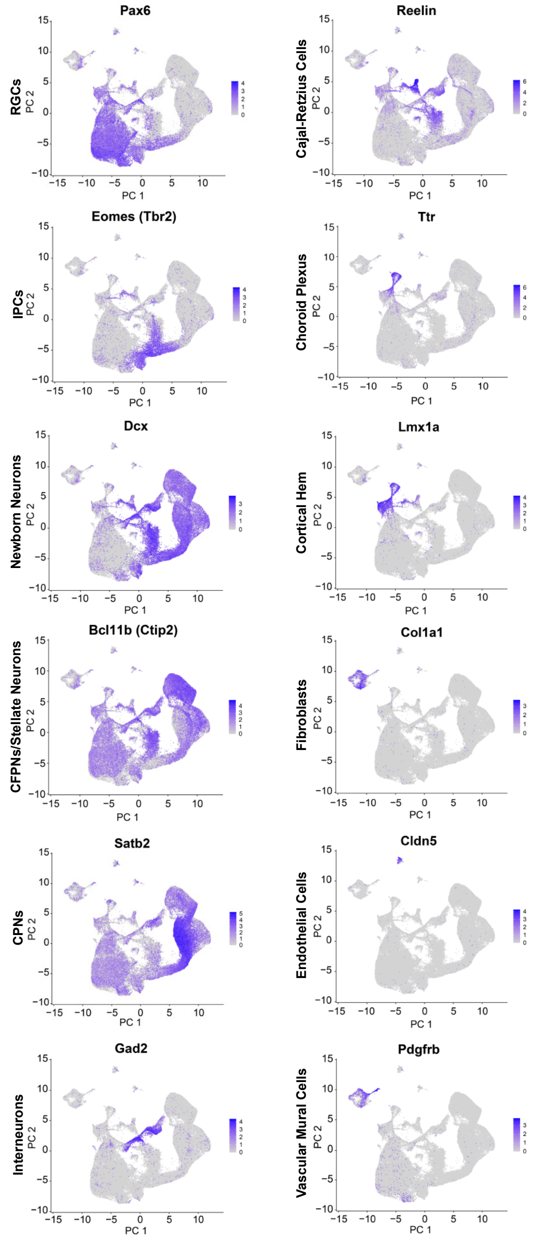
