## Supplemental Figure 3 for "*Loss of miR-9-2* Causes Cerebral Hemorrhage and Hydrocephalus by Widespread Disruption of Cell-Type-Specific Neurodevelopmental Gene Networks"

A

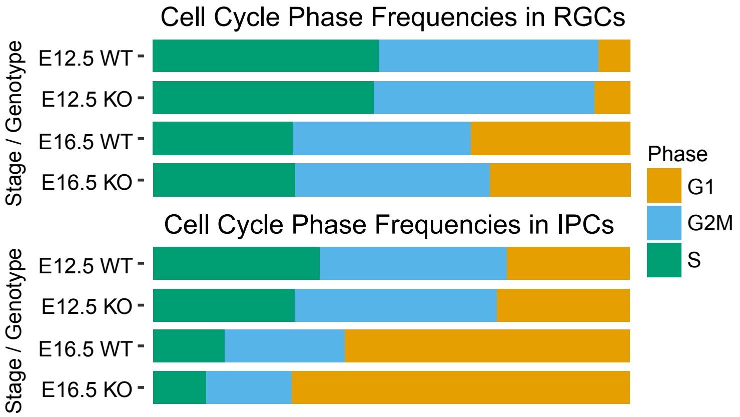

B

| Cell Type | Stage | Phase | logFC | logCPM | F | PValue | FDR |
|---|---|---|---|---|---|---|---|
| RGCs | E12 | G1 | 0.186 | 16.149 | 0.429 | 0.521 | 0.977 |
| RGCs | E12 | S | -0.035 | 18.842 | 0.031 | 0.863 | 0.977 |
| RGCs | E12 | G2M | 0.006 | 18.805 | 0.001 | 0.977 | 0.977 |
| RGCs | E16 | G1 | -0.216 | 18.183 | 0.357 | 0.567 | 0.894 |
| RGCs | E16 | G2M | 0.138 | 18.656 | 0.142 | 0.716 | 0.894 |
| RGCs | E16 | S | 0.028 | 18.164 | 0.019 | 0.894 | 0.894 |
| **Cell Type** | **Stage** | **Phase** | **logFC** | **logCPM** | **F** | **PValue** | **FDR** |
| IPCs | E12 | S | -0.165 | 18.308 | 0.709 | 0.414 | 0.819 |
| IPCs | E12 | G1 | 0.109 | 18.061 | 0.178 | 0.680 | 0.819 |
| IPCs | E12 | G2M | 0.059 | 18.631 | 0.054 | 0.819 | 0.819 |
| IPCs | E16 | S | -0.405 | 17.000 | 6.170 | 0.042 | 0.125 |
| IPCs | E16 | G1 | 0.244 | 19.290 | 2.097 | 0.191 | 0.286 |
| IPCs | E16 | G2M | -0.473 | 17.812 | 1.186 | 0.312 | 0.312 |

C

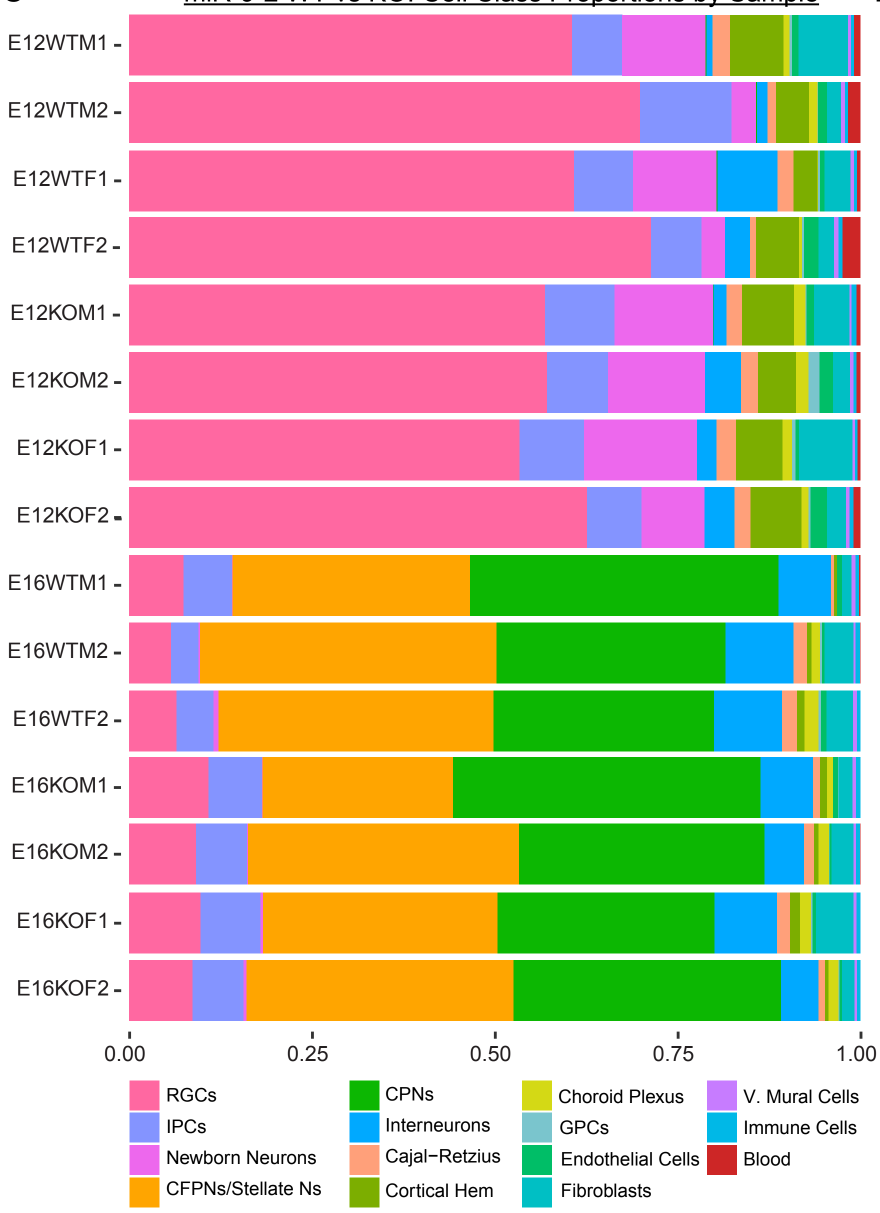

D

| E12.5 Differential Abundance of Cell Types by Genotype | | | | | |
|---|---|---|---|---|---|
| **Cell Type** | **logFC** | **logCPM** | **F** | **PValue** | **FDR** |
| RGCs | -0.192 | 19.232 | 1.193 | 0.305 | 0.522 |
| IPCs | -0.015 | 16.398 | 0.004 | 0.954 | 0.954 |
| Newborn Neurons | 0.786 | 16.615 | 2.704 | 0.138 | 0.448 |
| Interneurons | -0.054 | 15.074 | 0.006 | 0.939 | 0.954 |
| Cajal-Retzius Cells | 0.456 | 14.323 | 1.880 | 0.206 | 0.518 |
| Cortical Hem | 0.296 | 15.833 | 1.110 | 0.321 | 0.522 |
| Choroid Plexus | 1.106 | 13.387 | 4.318 | 0.071 | 0.440 |
| GPCs | 1.491 | 12.038 | 3.394 | 0.101 | 0.440 |
| Endothelial Cells | 0.294 | 13.725 | 0.346 | 0.572 | 0.827 |
| V. Mural Cells | -0.421 | 12.139 | 1.608 | 0.239 | 0.518 |
| Fibroblasts | 0.253 | 15.278 | 0.239 | 0.638 | 0.829 |
| Immune | 0.069 | 12.319 | 0.050 | 0.829 | 0.954 |
| Blood | -1.116 | 13.355 | 4.606 | 0.064 | 0.440 |
| **E16.5 Differential Abundance of Cell Types by Genotype** | | | | | |
| **Cell Type** | **logFC** | **logCPM** | **F** | **PValue** | **FDR** |
| RGCs | 0.552 | 16.349 | 12.180 | 0.009 | 0.107 |
| IPCs | 0.516 | 15.991 | 6.319 | 0.038 | 0.226 |
| CFPNs | -0.169 | 18.409 | 0.973 | 0.354 | 0.876 |
| CPNs | 0.033 | 18.425 | 0.032 | 0.863 | 0.941 |
| Interneurons | -0.380 | 16.186 | 2.976 | 0.125 | 0.499 |
| Cajal-Retzius Cells | -0.150 | 13.801 | 0.092 | 0.770 | 0.923 |
| Cortical Hem | 0.424 | 12.947 | 0.855 | 0.384 | 0.876 |
| Choroid Plexus | 0.336 | 13.597 | 0.214 | 0.657 | 0.876 |
| Endothelial Cells | -0.236 | 12.457 | 0.444 | 0.525 | 0.876 |
| V. Mural Cells | -0.187 | 12.024 | 0.301 | 0.599 | 0.876 |
| Fibroblasts | 0.021 | 14.881 | 0.002 | 0.967 | 0.967 |
| Immune | 0.135 | 12.494 | 0.336 | 0.579 | 0.876 |
